# Pheno-Morphological Characterization and Genetic Diversity Assessment of Grain and Vegetable Soybean (Glycine max. (L.) Merrill) Lines for Breeding Advancements

**DOI:** 10.1101/2024.12.01.626274

**Authors:** Madhumita Saha, Mohan Chavan, T Onkarappa, R L Ravikumar, Uddalak Das

**Affiliations:** Department of Plant Biotechnology, University of Agricultural Sciences, Bangalore, Bengaluru, Karnataka, India

**Keywords:** Phenotyping, edamame, vegetable soybean, grain soybean, genetic variability studies

## Abstract

This study presents for the first time, a thorough pheno-morphological characterization of ten soybean [*Glycine max* (L.) Merrill] lines (four grain type and six vegetable type). The vegetable soybean, also called as *Edamame*, and the grain type soybean, which underpin considerable genetic variation across traits such as flowering and pod-setting times, plant height, and seed size. Analyses of variance and clustering reveal obvious differentiation between vegetable and grain-type soybeans, particularly in traits such as height at the R7 stage, pod length, and branching patterns. The vegetable type exhibited faster plant development throughout reproductive stages and tended to be taller than the grain type. The study found characteristics, including test weight, plant height, and pods per plant, had higher phenotypic and genotypic coefficient of variation and, therefore, may serve as good candidates for selection breeding since these characteristics are based on a strong genetic constitution and to a lesser extent on environmental influences. The traits like number of pods per plant and test weight had high heritability and genetic advance and thus held great promise for improvement of productivity through selection. Important PCA explained traits were the plant height at the R7 stage, number of branches, and number of pods per cluster, which have high values for the breeding program. Hierarchical clustering also clumped the genotypes from other morphological characteristics which grouped them into clear distinct classification for the genetic diversity of this germplasm. In relation to this, one realizes that these soybean lines have genetic variability for significant breeding programs concerning increases in yield, adaptability, and resistance to stresses: biotic and abiotic.

## 1. Introduction

It is one of the legume crops widely grown in the world and has two types, namely, grain soybean and vegetable soybean. This also known as “Golden bean” and miracle crop of 20th century because of their multi-effects. The vegetable soybean, or edamame [*Glycine max* (L.) Merrill] is a species in the family Fabaceae (Singh, 2017). Vegetable soybeans were cultivated as far back as the 11th century in China (Hymowitz and Shurtleff, 2005). Vegetable soybean, though not commonly cultivated in India, has much promise in many regions with favourable agricultural conditions. Vegetable soybean, which boasts the greatest protein yield per unit area, has much potential to improve nutritional deficiency significantly. *Glycine max* (L.) Merrill is the large-seeded variety of soybean and is most commonly harvested at the R6 growth stage when the pods are green. In India, the productivity of vegetable soybean lags the world average for many reasons: low genetic diversity, narrow genetic base of vegetable soybean varieties, a short growing period dictated by India’s latitudinal position, and stagnant genetic yield potential (Satyavathi *et al*., 2003). Plant morphological characters form the basis for characterizing the genotypes of soybeans, including grain and vegetable varieties. These include morphological characters like anthocyanin pigmentation on the hypocotyl, leaf shape, growth habit, flower color, presence, and color of pod pubescence, pod color, seed size, seed, and cotyledon color. These characters have been identified as distinguishing characters to characterize, identify, and verify soybean genotypes. The knowledge of the inheritance of qualitative traits provides an aid to describe new genes, construct linkage maps, and formulate marker assisted selection methods. Because of its abundance in oil, protein, and medicinal components, soybeans have been widely used as raw materials for various industries. The germplasm review for valuable traits requires that the knowledge on genetic diversity be followed up continuously. Before that, this evaluation was based only on the morphological data available. In addition to reflecting the genetic makeup of the crop, the morphological characters and markers provide evidence for how the genotype interacts with its environment of expression.

Shurtleff and Aoyagi (2001); Shurtleff and Lumpkin (2001) report that green beans are eaten directly from the pods. Vegetable soybean, also known as edamame, makes up a small portion of the total world soybean production at around 2 percent. Vegetable soybean seeds are sweeter and softer compared to grain soybean seeds (Shanmugasundaram and Yan, 2010) as well as considerably larger in size (>250 mg/seed or >30 g/100 seeds dry weight) compared to grain soybean seeds (Carson, 2010). Ravikumar and Narayanaswamy (1999) have concluded that seed coat color, hilum color, seed shape, and seed size are stable for varietal identification. Clarke and Wiseman (2000) noted that there was a large genetic variability in the vegetable soybean accessions. They also observed remarkable differences between the vegetable soybean types concerning T.S.S. (%), green pod production per plant, test weight 100 of fresh seed and pod length. There is room for selection for these traits among the vegetable soybean genotypes due to the high heritability combined with high GAM for plant height, green pod yield per plant, test weight 100 of fresh seed, T.S.S. (%), and pod length. Since there was little genetic variation found for pod width, both heritability and GAM were low. Kumar *et al*., 2015, estimated forty genotypes of soybeans for genetic factors and agro-morphological features. For any of these characteristics, correlation and path coefficients were generated. Significant differences between all the genotypes were found for each character when mean performance and analysis of variance for yield and its components were conducted. High significant correlation was also identified between yield and the qualities that make it up. According to a route analysis, the hundred seed weight affected the yield the most. Genetic architecture is a product of vast natural selection among a population. A population that inhabits a range of different habitats may have a highly heterogeneous genetic makeup. Multivariate analysis, D^2^ analysis created by Mahalanobis in 1936, has been found to be an effective one among the several statistical techniques developed to determine divergence across populations (Rao, 1960). Divergence assessment is useful for parent selection in a given breeding objectives as it is well known that the genetic divergence of the parents in a hybridization process contributes significantly to the success of a breeding program. Summary of D^2^ analysis literature. This is a brief summary of the D^2^ analysis literature.

## 2. Materials and Method

### 2.1. Experimental details

The ten lines were treated as variables with three replications and four were considered as checks when the field was prepared using the RCBD (Randomized Complete Block Design) approach. The plants were sown at a spacing of 30 cm × 35 cm (Row to Row and Plant to Plant).

### 2.2. Plant material

Soybean lines were collected from AICRP Soybean, GKVK campus, UAS Bengaluru. The total number of genotypes used were ten, out of which five were vegetable type soybeans and five were checks (1 vegetable type and 4 grain type). The check varieties used were Karune (Vegetable type) and RKS 18, KBS 23, EC 0916018 and EC 76758 (Grain type). The experimental material comprised of five vegetable germplasm soybean lines (EC916019, EC916029, EC892880, EC892882, and EC892808), along with check variety Karune, were chosen as vegetable check for the experiment. These soybean lines were experimented with in addition to two grain-type soybeans, KBS-18 and RKS-23 and two grain-type bold-seeded soybeans, EC 0916018 and EC 76758 and considered as the grain check line (**Figure 1**).

**Figure 1:**
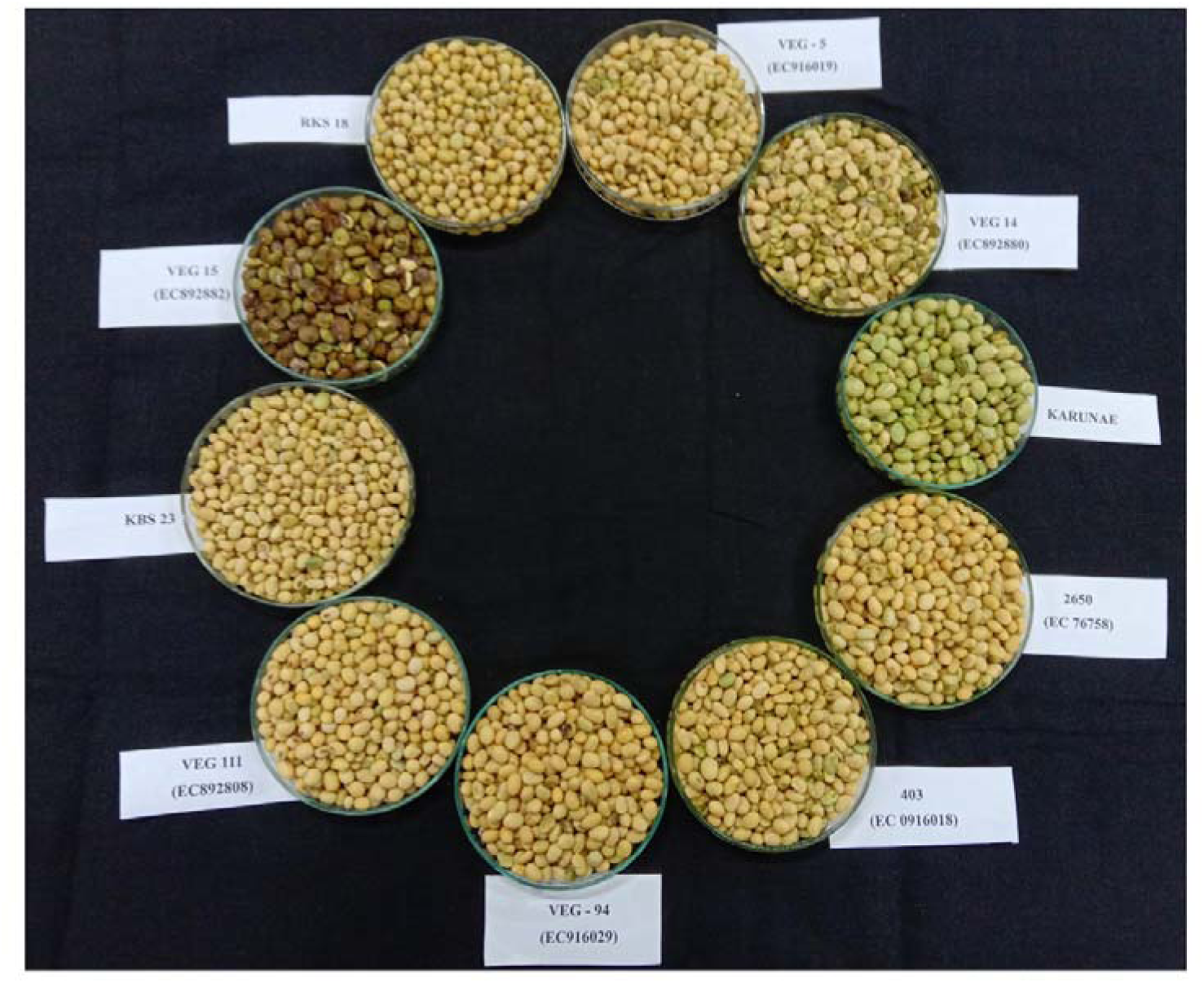
All ten soybean lines used in the study including both six vegetable types and four grain types.

### 2.3. Pheno-morphological characterization of five plants per genotypes per replication

Quantitative traits like days to 50% flowering, i.e., the days taken to attain 50 percent of the plants to flower was measured starting from the day of sowing; days for pod setting, in which the count of days a certain accession of the plant was grown was followed from the moment of sowing until the first pod of its germination was recorded on the plants; number of days taken to attain R7 stage, in which, five representative random plants were chosen and tagged, these plants were used for taking observation from the R7 and R8 stage of growth; plant height at R7 stage (cm), i.e., the heights of the base to the tip of five plants of each genotype were measured in centimetres using scale and noted down. Afterwards, the average plant height that pertains to each genotype was obtained; number of branches per plant, in which the number of branches per plant was counted from five selected plants at harvest, and the mean value were calculated for each genotype; number of pods per plant, where, the number of pods per plant was manually counted at R4 stage from five selected plants. The mean number of pods per plant was then calculated based on these counts; length of pod (cm), i.e., the length of pods for each genotype was measured at R6 stage using a vernier calliper from the base of the sepals to the tip of the pod. Five plants were taken into observation per genotype per replication and the average pod length was calculated; number of pods per cluster, where, the total number of pods per cluster from randomly selected five clusters were meticulously counted and accurately recorded during the harvest process; number of seeds per pod, i.e., the number of seeds per pod at R7 stage from five randomly selected pods were manually counted and the mean number of seeds per pod was determined by calculating the average of these counts; test seed weight (grams), in which the weight of the hundred seeds were measured and recorded for selected genotypes, normally if the seeds are small, 1000 seed weight and if the seeds are bold seeded, 100 seed is considered as test weight of the sample; seed size (mm), where randomly selected five dried soybean plant’s seeds were individually measured using a vernier calliper to determine their size. The measurements were averaged to obtain the average size of the seeds. Qualitative traits like plant growth habit, categorized as determinate or indeterminate based on observed characteristics. Determinate plants typically stop growing once they reach a certain size or maturity, while indeterminate plants continue to grow and produce indefinitely under favourable conditions. Determinate plants are compact and bushy, while indeterminate plants are taller and more open; corolla colour, i.e., the colour of the flower was documented as either white or purple; seed shape, i.e., the mature seed shape is categorized based on its appearance as either semi-flat or globular; seed coat colour, where the seed coat colour at R7 stage was observed under natural light and classified into two groups: green and yellow; pubescence density, i.e., the pubescence on leaf, stem, and pod was visually examined and recorded, and genotypes were categorized based on level of presence or absence; mature pod colour, where in each genotype, the mature pods were visually observed to categorize them based on colour as either brown or olive green, leaflet shape, where the genotypes were categorized into four groups based on the shape of their lateral leaflet: lanceolate, triangular, pointed ovate, and rounded ovate. Each shape classification likely corresponds to distinct genetic traits or phenotypic characteristics within the studied population.

### 2.4. Reproductive growth stages of soybean lines

By counting the days, the number of days needed to reach each reproductive stage has been determined. Days were counted until the R6 stages since it is advised to harvest vegetable soybeans at this stage since pod development would have reached its physiological maturity. R1 stage, when the crop begins to flower, it reaches the R1 stage. When a plant has at least one flower on any given node, flowering has begun. From each genotype, five plants were chosen at random, and the number of days it took for them to begin flowering was noted; R2 stage, when every plant in the population reaches full flowering, the crop reaches the R2 stage. When one of the two topmost nodes has an open flower, full flowering has occurred. To determine the R2 stage for the plant population, five randomly selected plants were counted down to the end of the day; R3 stage, upon the onset of pod development within the plant population, the crop reaches the R3 stage. At one of the top four nodes, 3/16- inch (5 mm) long pods begin to grow. The number of days it took for five randomly selected plants to reach the R3 stage was calculated; R4 stage, when the crop’s pod growth is seized, it reaches the R4 stage. It reaches one of the four highest nodes, where its pod size reaches ¾ inch (2 cm). Five plants were selected at random, and the number of days it took for the pods to fully grow has been noted; R5 stage, when seed development commences, the crop reaches the R5 stage. At one of the four topmost nodes on the main stem, the pod begins to generate seeds that are 1/8 inch (3 mm) long; R6 stage, when the entire seed population has developed, the crop reaches the R6 stage. One of the four topmost nodes on the main stem’s food capacity is filled by the green seeds found in the pods; R7 stage, when one mature normal pod (brown or tan) is present on the main stem. In seeds, dry matter is peaking. The green is fading and the pods and seeds look yellow. When the seeds reach physical maturity, 60% of them are wet. Stress doesn’t really matter unless the pods break or fall to the ground.

### 2.5. Statistical analysis

The statistical procedures were performed in R-studio software.

#### 2.5.1. Analysis of variance

Analysis of variance (ANOVA) was conducted using an RCBD Excel sheet to assess variability among ten genotypes across measurable traits. Data analysis was performed in R Studio following the standard RCBD procedure. Treatment differences were deemed significant if the calculated F-value exceeded the tabulated F-value at a 5% significance level (**Table 1**). Variance and covariance analyses for individual traits and trait pairs were based on plot mean values (Panse and Sukhatme, 1954).

**Table 1:** Structure of ANOVA as per randomized block design quantitative characters in vegetable soybean genotypes.

| Source of variation | Degrees of freedom | Mean sum of squares | Expected mean sum of squares |
| --- | --- | --- | --- |
| Replication | (r - 1) | $\mu_{RSS}$ | $\sigma^2_e + \sigma^2_r$ |
| Genotype | (g - 1) | $\mu_{GSS}$ | $\sigma^2_e + r \cdot \sigma^2_g$ |
| Error | (r - 1) (g - 1) | $\mu_{ESS}$ | $\sigma^2_e$ |
| Total | (rg - 1) |  |  |

where, r = number of replications; g = number of genotypes; $\sigma^2_e$ = Variance due to error; $\sigma^2_r$ = Variance due to replication; $\sigma^2_g$ = Variance due to genotype; $\mu_{RSS}$ , $\mu_{GSS}$ , $\mu_{ESS}$ = Replication, genotype and error mean sum of squares respectively

where, r = number of replications; g = number of genotypes; σ^2^e = Variance due to error; σ^2^r = Variance due to replication; σ^2^g = Variance due to genotype; µ_RSS,_ µ_GSS,_ µ_ESS_ = Replication, genotype and error mean sum of squares respectively

#### 2.5.2. Variability parameters

To assess genetic variability among genotypes, phenotypic data for quantitative traits were analyzed using basic statistical measures, including mean, range, variance, and standard error. Genetic parameters like genotypic coefficient of variance (GCV) (Burton, 1952) and broad-sense heritability [h² (bs)] (Lush, 1945) were calculated based on mean trait values for each genotype.

Mean:

Mean is ratio of sum of all the observations divided by total number of observations.

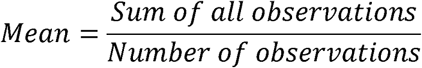

Range:

Range is the difference between highest and lowest values of the observation for a trait.

Standard error:

Standard error is the measure of uncontrolled variations in the sample. it is calculated by dividing standard deviation by the square root of number of observations in sample. It is denoted by SE.

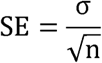

Where, σ = standard deviation; n = Number of observations Genotypic Co-efficient of Variation (GCV)

Genotypic co-efficient of variation for all the characters were estimated using the formula as suggested by the Burton (1952).

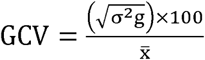

Where, σ^2^g = Genotypic variance; XD= Population mean

The GCV was classified into low (0 – 10 %), moderate (10 % - 20 %) and high (> 20 %) (Robinson *et al.,* 1949).

Phenotypic Co-efficient of Variation (PCV) The phenotypic coefficient of variation (Burton, 1952) was calculated from the following formula,

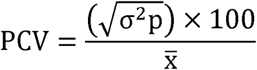

Where, σ^2^p = Phenotypic variance; XD= Population mean

#### 2.5.3. Principal Component Analysis

Principal Component Analysis (PCA) is one of the most widely used multivariate statistical methods across scientific fields. Although its modern form was developed by Hotelling in 1933, PCA’s origins trace back to Pearson in 1901. The technique analyzes data tables with observations across multiple dependent variables, aiming to distill essential data into new, orthogonal components. By mapping these components, PCA visually highlights patterns of similarity among variables and observations (Jolliffe, 2002; Jackson, 1991).

Using a data-driven modeling approach called principal component analysis (PCA), a group of correlated variables can be reduced to a smaller set of new variables that are uncorrelated and retain most of the original data. Let X be the input dataset, with m- dimensional input sequences in each column. Furthermore, remember that each function in the dataset has an average of zero [E(X) = 0]. An original data matrix can generally be given with n samples and m variables as follows:

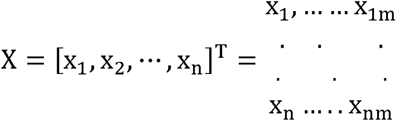

Through the use of PCA, weather variable data and performance parameter input can be transformed into a new event space while maintaining as much of the original data as possible. In order to do this, the input data sets are searched for the directions of maximum variance, and then they are projected to a new subspace with dimensions that are either equal to or less than those of the original space. Thus, an orthonormal transformation Z may be used to transfer X to new space T as follows:

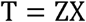

The X-matrix elements are combined linearly to create the orthonormal vectors that make up the T-matrix of scores, which represents the relationships between the samples. The T covariance matrix is:

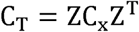

X’s covariance matrix is denoted by C_X_. The eigenvalue equation can be used to generate the loading matrix Z, which is written as:

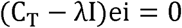

The covariances pairwise between the several input variables are stored in the covariance matrix. The covariance matrix’s eigenvectors and eigenvalues are then decomposed, and the related eigenvectors characterize the new orthogonal components—also referred to as “principal components”—while the corresponding eigenvalues will decide how big these components are. The first principal component will have the biggest potential variance (i.e., information), followed by the second principle component with the second largest variance, and so on, once the eigenvalues and their matching eigenvectors have been ranked in descending order. It should be mentioned that even in cases when the input variables exhibit correlation because to the orthogonality of the decomposed eigenvectors, the output principal components do not exhibit correlation with one another.

#### 2.5.4. Heritability (h^2^)

Broad sense heritability was estimated (Lush 1949) by the formula suggested by Johnson *et al*. (1955), Hanson *et al*. (1956).

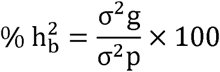

Where, h^2^ _b_ = Heritability; σ^2^_g_ = Genotypic variance; σ^2^ = Phenotypic variance The heritability was categorized as Low (5-10 %); Moderate (10-30 %) and High (>30 %) Robinson (1949).

#### 2.5.5. Genetic Advance

Genetic Advance (GA) and percentage of the mean (GAM) assuming selection of superior 5% of the genotypes was estimated in accordance with the methods illustrated by Johnson *et al*. (1955) as:

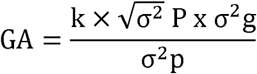

Where, GA = Expected genetic advance; k = Standardized selection differential at 5% selection intensity (K = 2.063); σ^2^p = Phenotypic variance; σ^2^g = Genotypic variance

2.5.6. **Genetic advance as percentage of mean (GAM)**

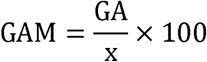

Where, GAM = Genetic advance as percentage of mean; GA = Expected genetic advance; x = Grand mean of a character The GAM was classified into low (0 – 10 %), moderate (10 % - 20 %) and high (> 20 %) (Robinson *et al.,* 1949).

#### 2.5.7. Intra and inter cluster distances

The intra and inter-cluster distance (D) were calculated by taking the square roots of respective D^2^ values between genotypes within a particular cluster, and between genotypes belonging to separate cluster combinations, respectively.

The genotypic divergence between populations was evaluated using the Mahalanobis D^2^ statistic (Mahalanobis, 1936). The generalized distance between any two populations is determined by formula:

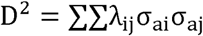

Where, D^2^ = Square o generalized distance; λ_ij_ = Reciprocal of the common dispersal matrix; σ_ai_ = ( μ_i1_ - μ_i2_); σ_aj_ = ( μ_j1_ - μ_j2_); μ = General mean The computing process was made simpler by transforming the initial correlated unstandardized character mean (Xs) into standardized uncorrelated variable (Ys), as the computation formula necessitates the inversion of a higher order determinant. The sum of squares of the differences between pairs of corresponding uncorrelated (gs) values of any two uncorrelated genotypes of D^2^ value was used to calculate the D^2^ values.

Toucher’s approach, as explained by Rao (1952), was used to cluster all n (n-1) / 2D^2^ values. With the following formula, the intra cluster distances were computed (Singh and Choudhary, 1997).

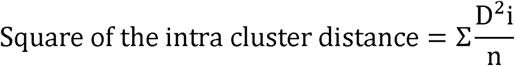

Where, Σ D^2^i is the number of possible combinations (n) is equal to the total of the distances between all conceivable combinations of the elements that make up a cluster.

The following formula was used to compute the inter-cluster distances, as stated by Singh and Choudhary (1997):

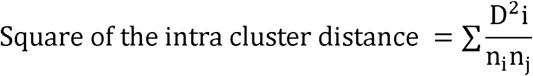

In the clusters under investigation, ΣD^2^_i_ represents the total distance between all conceivable combinations (n_i_ n_j_) of the entries. The number of items in Cluster I is denoted by n_i_, and in Cluster J by n_j_. This method’s criterion for clustering was that any two genotypes that belong to the same cluster should, on average, have smaller D^2^ values than genotypes that belong to two separate clusters.

## 3. Results and discussion

### 3.1. Pheno-morphological characterization of vegetable soybean lines

Analysis of variance revealed highly significant variation across vegetable soybean lines, indicating sufficient diversity in the germplasm. Recorded traits included days to 50% flowering, pod setting, days to R7 stage, plant height at R7, branch and pod numbers, pod length, seeds per pod, test seed weight, seed size, growth habit, corolla and seed coat colors, pubescence density, mature pod color, and leaflet shape (**Table 2**).

**Table 2:** Mean values of quantitative traits of the ten soybean genotypes.

| Soybean lines | DTF | DPS | DTRR7 | PHR7 | NBPP | NPPP | PL | NPPC | NSPP | TW | SS |
| --- | --- | --- | --- | --- | --- | --- | --- | --- | --- | --- | --- |
| **EC916019** | 42.56 ± 0.2 | 67.58 ± 0.5 | 77.66 ± 0.5 | 62.54 ± 4.9 | 14.00 ± 0.9 | 32.33 ± 2.3 | 5.44 ± 0.20 | 6.86 ± 0.6 | 2.98 ± 0.2 | 20.44 ± 0.10 | 8.01 ± 0.40 |
| **EC916029** | 43.00 ± 0.4 | 66.06 ± 0.5 | 77.26 ± 0.5 | 65.24 ± 1.6 | 13.73 ± 0.7 | 33.33 ± 2.8 | 4.99 ± 0.28 | 6.76 ± 0.8 | 2.73 ± 0.2 | 21.47 ± 0.10 | 8.08 ± 0.44 |
| **EC892880** | 41.66 ± 0.3 | 67.40 ± 0.6 | 77.80 ± 0.4 | 64.08 ± 1.1 | 12.09 ± 0.9 | 35.60 ± 3.9 | 5.53 ± 0.40 | 6.93 ± 0.9 | 2.79 ± 0.3 | 19.26 ± 0.20 | 8.02 ± 0.50 |
| **EC892882** | 40.00 ± 0.2 | 66.86 ± 0.4 | 77.40 ± 0.7 | 50.51 ± 1.8 | 14.60 ± 0.9 | 33.73 ± 2.6 | 5.63 ± 0.23 | 6.56 ± 0.6 | 2.81 ± 0.3 | 21.72 ± 0.10 | 9.02 ± 0.48 |
| **EC892808** | 41.00 ± 0.5 | 67.90 ± 0.4 | 78.12 ± 0.4 | 55.24 ± 1.5 | 13.20 ± 1.3 | 38.53 ± 2.1 | 5.54 ± 0.19 | 6.66 ± 0.8 | 2.66 ± 0.3 | 22.28 ± 0.20 | 7.42 ± 0.80 |
| **Karune** | 40.63 ± 0.3 | 67.86 ± 0.6 | 78.01 ± 0.8 | 51.22 ± 2.5 | 12.53 ± 1.8 | 39.60 ± 1.7 | 5.44 ± 0.22 | 6.46 ± 1.1 | 2.86 ± 0.2 | 20.62 ± 0.09 | 9.24 ± 0.56 |
| **RKS 18** | 43.00 ± 0.3 | 69.93 ± 0.5 | 78.91 ± 0.9 | 34.13 ± 1.8 | 11.66 ± 1.5 | 23.86 ± 1.5 | 5.32 ± 0.27 | 6.01 ± 0.8 | 2.36 ± 0.6 | 15.46 ± 0.12 | 6.03 ± 0.72 |
| **KBS 23** | 44.00 ± 0.2 | 71.00 ± 0.9 | 80.40 ± 0.6 | 31.07 ± 1.4 | 10.73 ± 1.9 | 27.53 ± 1.4 | 5.34 ± 0.19 | 6.26 ± 0.6 | 2.18 ± 0.1 | 14.49 ± 0.11 | 6.33 ± 0.43 |
| **EC76758** | 45.00 ± 0.4 | 70.50 ± 1.4 | 79.40 ± 0.6 | 42.11 ± 1.5 | 10.86 ± 1.5 | 25.20 ± 1.3 | 4.77 ± 0.13 | 5.26 ± 0.5 | 2.56 ± 0.2 | 24.72 ± 0.03 | 10.64 ± 0.6 |
| **EC0916018** | 44.00 ± 0.4 | 69.73 ± 0.6 | 78.86 ± 0.7 | 41.22 ± 1.3 | 10.66 ± 1.5 | 28.53 ± 1.1 | 4.69 ± 0.23 | 5.60 ± 0.7 | 2.43 ± 0.3 | 23.32 ± 0.06 | 9.89 ± 0.35 |
| **Mean** | **42.48** | **68.48** | **78.38** | **49.73** | **12.40** | **31.82** | **5.27** | **6.34** | **2.63** | **20.38** | **8.27** |
| **Minimum** | **40.00** | **66.06** | **77.26** | **31.07** | **10.66** | **23.86** | **4.69** | **5.26** | **2.18** | **14.49** | **6.03** |
| **Maximum** | **45.00** | **71.00** | **80.40** | **65.24** | **14.60** | **39.60** | **5.63** | **6.93** | **2.98** | **24.72** | **10.64** |
| **CV (%)** | **1.79** | **0.87** | **0.54** | **3.02** | **14.00** | **4.19** | **3.29** | **14.57** | **6.87** | **0.65** | **4.85** |
| **CD (5%)** | **1.40** | **1.10** | **0.79** | **2.77** | **3.20** | **2.46** | **0.32** | **1.70** | **0.33** | **0.24** | **0.74** |
| **CD (1%)** | **0.42** | **1.40** | **1.00** | **3.53** | **4.08** | **3.13** | **0.40** | **2.16** | **0.42** | **0.31** | **0.94** |

**Note:** **DPS:** Days for pod setting; **DTRR7:** Days Taken to Reach R7 stage; **DTF:** Days To 50% Flowering; **TW:** Test Weight; **NBPP:** Number of Branches Per Plant; **NPPC:** Number of Pods Per Cluster; **NPPP:** Number of Pods Per Plant; **NSPP:** Number of Seeds Per Pod; **PHR7:** Plant Height at R7 stage; **PL:** Pod Length; **SS:** Seed Size

#### 3.1.1. Days to 50% flowering

Among the observed vegetable lines, flowering duration showed notable variation, with an average of 42.48 days and a range from 40.00 to 45.00 days. The fastest to flower was variety EC892882 at 40 days, while grain type EC76758 took the longest at 45 days. Other varieties, including Karune, RKS 18, KBS 23, and EC0916018, reached flowering between 40.63 and 45.00 days (**Table 2**). This aligns with findings by Khanande *et al*. (2016), who observed longer flowering durations in grain-type soybean genotypes like TAMS-98-21 (46.70 days) and shorter durations in vegetable genotypes like AGS-450 (41.56 days).

#### 3.1.2. Days for pod setting

In the ten soybean genotypes studied, days to pod setting ranged from 66.06 to 71.00 days, with an average of 68.48 days. The vegetable type genotype EC916029 set pods the earliest (66.06 days), while grain type KBS-23 took the longest (71.00 days). Most vegetable types (EC916019, EC916029, EC892880, EC892882, and Karune) had similar pod-setting times (around 67 days), whereas grain types (RKS-18, KBS-23, EC76758, and EC0916018) averaged closer to 70 days (**Table 2**). These findings align with Zhang and Boahen (2007), who reported similar trends.

#### 3.1.3. Number of days taken to attain R7 stage

Vegetable-type soybeans showed a distinct variation in days to reach the R7 stage, averaging 78.38 days with a range from 77.26 to 80.40 days. The earliest line was EC916029 (77.26 days), while the grain-type line KBS-23 was the latest (80.40 days). Most vegetable-type lines reached R7 earlier than grain types (**Table 2**), consistent with findings by Zhang and Boahen (2007), where vegetable types reached R7 in 79.35 days compared to 82.6 days for grain types.

#### 3.1.4. Plant height at R7 stage (cm)

At the R7 stage, plant height across the population varied significantly, averaging

49.73 cm, with a range of 31.07 cm to 65.24 cm. The tallest line, EC 916029 (65.24 cm), was a vegetable soybean, while the shortest, KBS-23 (31.07 cm), was a grain type. Vegetable soybean lines showed greater heights (50.51 cm to 65.24 cm) compared to grain types (31.07 cm to 42.11 cm) (**Table 2**), aligning with findings by Khanande *et al*. (2016) for vegetable (63.04 cm) and grain type (54.12 cm) soybean lines.

#### 3.1.5. Number of branches per plant

Soybean branch counts varied significantly, averaging 12.4 branches per plant, with a range of 10.0 to 14.0. The vegetable type EC892882 had the highest branch count (14.6), while the grain type EC0916018 had the lowest (10.66) (**Table 2**). Similarly, Zhang and Boahen (2007) reported branch averages of 9.2 for vegetable soybeans and 8.0 for grain types.

#### 3.1.6. Number of pods per plant

The number of pods per branch in various seeded soybean lines ranged from 23.86 to 39.60, with a mean of 31.82. Karune recorded the highest at 39.60 pods per branch, while RKS-18 had the lowest at 23.86 pods per branch (**Table 2**). Zhang and Boahen (2007) reported similar findings, with the grain type averaging 14.4 pods per plant and the vegetable type 16.4 pods per plant, showing a higher pod count in the vegetable variety.

#### 3.1.7. Length of pod (cm)

A study of ten soybean genotypes revealed that vegetable-type soybean lines typically had slightly longer pods than grain-type lines, with pod lengths ranging from 4.69 cm in the grain-type line EC 0916018 to 5.63 cm in the vegetable-type line EC 892882, averaging 5.27 cm. Despite these differences, vegetable soybean lines and grain-type check varieties showed similar pod lengths. Bold-seeded grain-type lines like EC 76758 and EC 0916018 had shorter pods, measuring 4.77 cm and 4.69 cm, respectively (**Table 2**). This aligns with Khanande *et al*. (2016), where vegetable-type AGS-450 had 2.90 cm pods and grain-type IC-16815 had 5.40 cm pods.

#### 3.1.8. Number of pods per cluster

Among the vegetable lines tested, there was little variation in the number of pods per cluster, with an average of 6.34, ranging from 5.26 to 6.93. The lowest count (5.26 pods) was observed in the bold-seeded grain type soybean line EC76758, while the highest (6.93 pods) was found in the vegetable type soybean line EC892880 (**Table 2**). These results are consistent with those of Shilpashree *et al*. (2021), who reported 5.63 pods per cluster for vegetable types and 6.21 for grain types.

#### 3.1.9. Number of seeds per pod

The number of seeds per pod did not show significant variation among different soybean varieties, ranging from 2.18 to 2.98 on average. Vegetable soybean lines (EC916019, EC916029, EC892880, EC892882, EC892808, and Karune) had similar seed counts per pod (2.98 to 2.66), as did the grain-type check varieties (RKS 18, KBS 23, EC 76758, EC 0916018, and Karune) with counts ranging from 2.36 to 2.18 (**Table 2**). These findings align with Zhang and Boahen (2007), who reported similar seed counts for grain and vegetable types.

#### 3.1.10. Test seed weight (grams)

The weight of 100 seeds varied significantly across the tested lines, ranging from 14.49 g to 24.72 g, with an overall mean of 20.38 g. Among the vegetable-type soybean lines, EC892808, EC892882, EC916029, EC916019, and EC892880 had average weights of 22.28 g, 21.72 g, 21.47 g, 20.44 g, and 19.26 g, respectively. The grain-type line KBS-23 had the lowest test weight at 14.49 g, while EC 76758, a bold-seeded variety, recorded the highest at 24.72 g (**Table 2**). Similar findings were reported by Shilpashree *et al*. (2021), noting that vegetable soybeans had larger seeds with an average weight of 24.40 g, compared to 13.9 g for grain types.

#### 3.1.11. Seed size (mm)

The seed size of various soybean lines showed significant variation, with a mean of 8.27 cm, ranging from 6.03 cm to 10.64 cm. The largest seed size was observed in the bold- seeded grain type check variety (10.64 cm), while the smallest was in RKS-18 (6.03 cm). Vegetable-type soybean lines had moderate seed sizes, ranging from 7.42 cm (EC892808) to 9.24 cm (Karune) (**Table 2**). This aligns with the findings of Zhang and Boahen (2007), where vegetable-type soybeans (9.64 cm) had slightly larger seeds compared to grain-type soybeans (5.94 cm).

#### 3.1.12. Plant growth habit

All ten soybean genotypes, including the check varieties, exhibited an indeterminate growth habit (**Table 3**). This finding aligns with the results of Shilpashree *et al*. (2021), who observed the same growth habit in all vegetable-type soybeans.

**Table 3:** Comparative study of qualitative traits in vegetable type to grain type soybean lines.

| Sl. No. | Genotypes | Traits |  |
| --- | --- | --- | --- |
| 1. | EC916019, EC916029, EC892880, EC892882, EC892808, Karune, RKS 18, KBS 23, EC 76758, EC 0916018. | Plant growth habit | Indeterminate |
|  | No genotypes were recorded determinant type growth habit |  | Determinate |
| 2. | EC916029, EC892880, EC892882, EC892808, Karune, EC916019 | Corolla colour | White |
|  | EC 76758, RKS 18, KBS 23, EC 0916018. |  | Purple |
| 3. | EC916019, EC916029, EC892808, Karune, EC 76758, EC 0916018. | Seed shape | Globular |
|  | RKS 18, KBS 23, EC892882, EC892880. |  | Semi-flat |
| 4. | EC916019, EC916029, EC892880, EC892808, RKS 18, KBS 23, EC 76758, EC 0916018. | Seed coat colour | Brown |
|  | Karune. |  | Green |
|  | EC892882. |  | Olive green |
| 5. | EC 76758, RKS 18, KBS 23, EC 0916018. | Pubescence of leaves, stem and pod | High |
|  | EC916019, EC892880 |  | Moderate |
|  | EC892882, EC892808, Karune, EC916029, EC 76758. |  | Low |
| 6. | EC916019, EC916029, EC892880, EC892808, RKS 18, KBS 23, EC 76758, EC 0916018. | Mature pod colour | Tan |
|  | EC892882, Karune. |  | Brown |
| 7. | EC916019, EC 76758. | Leaflet shape | Lanceolate |
|  | EC916029, EC892880, EC892882, EC892808, EC 0916018. |  | Pointed |
|  | Karune, RKS 18, KBS 23. |  | Round |

#### 3.1.13. Corolla colour

Out of ten soybean genotypes, including the check varieties, six exhibited white corolla color, while four had purple corolla color (**Table 3**). Nzaranyimana (2017) also reported white corolla color in vegetable soybean.

#### 3.1.14. Seed shape

Eight genotypes exhibited globular seed shapes, while two had semi-flat seeds (**Table 3**). This finding aligns with Khanande *et al*. (2016), who also observed a predominance of globular seeds.

#### 3.1.15. Seed coat colour

The predominant seed coat color across the ten genotypes was brown, observed in eight genotypes, followed by green in one and olive green in another (**Table 3**). Khanande *et al*. (2016) also reported that most vegetable-type soybeans have a brown seed coat.

#### 3.1.16. Pubescence density

Pubescence density on the leaves, stem, and pod was categorized as high, moderate, or low across all ten genotypes, including the check varieties. Four genotypes showed high pubescence, two had moderate, and four exhibited low pubescence (**Table 3**). Vegetable-type soybeans, as noted by Khanande *et al*. (2016), generally exhibit pubescence.

#### 3.1.17. Mature pod colour

Out of the ten genotypes, including the check varieties, eight displayed tan pod color, while two showed brown (**Table 3**). Khanande *et al*. (2016) also reported the dominance of ten pod colors in vegetable-type soybeans.

#### 3.1.18. Leaflet shape

In a study of ten soybean genotypes, including check varieties, the most common leaflet shape was pointed, found in five genotypes. Round-shaped leaves were observed in three genotypes, while lanceolate leaves appeared in two (**Table 3**). Shilpashree *et al*. (2021) reported an equal distribution of all three leaflet shapes in vegetable-type soybeans.

For qualitative traits, no significant differences were inferred between the vegetable type and grain type soybean lines, except for the difference in their pubescence density and corolla colour (**Figure 2**).

**Figure 2:**
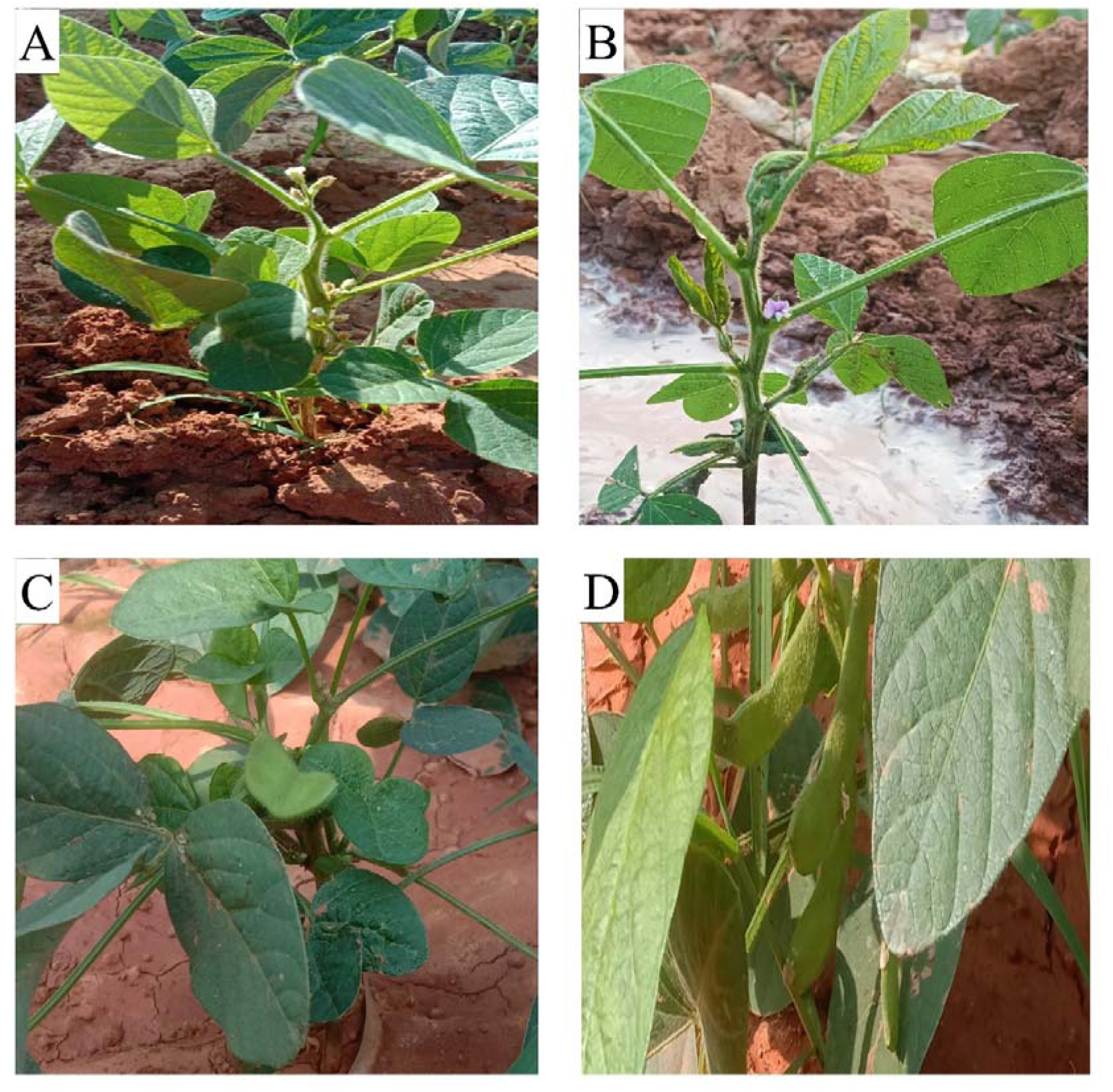
**A.** Vegetable type soybean flower**; B.** Grain type soybean flower ; **C.** Vegetable type soybean pods, **D.** Grain type soybean pods

### 3.2. Hierarchical clustering using pheno-morphological traits

Hierarchical clustering was performed on ten genotypes based on ten quantitative traits, using squared Euclidean distance and Ward’s method. The dendrogram, created through UGPMA clustering, grouped the genotypes into two main clusters. One cluster contained sub-clusters of vegetable-type soybeans, such as EC916019, EC916029, and EC892880, while another sub-cluster included EC892882, EC892808, and Karune (**Figure 3**). The second major cluster was divided into two sub-clusters: one with RKS-18 and KBS- 23 (grain-type soybeans), and another with EC76758 and EC0916018 (bold-seeded grain types).

**Figure 3:**
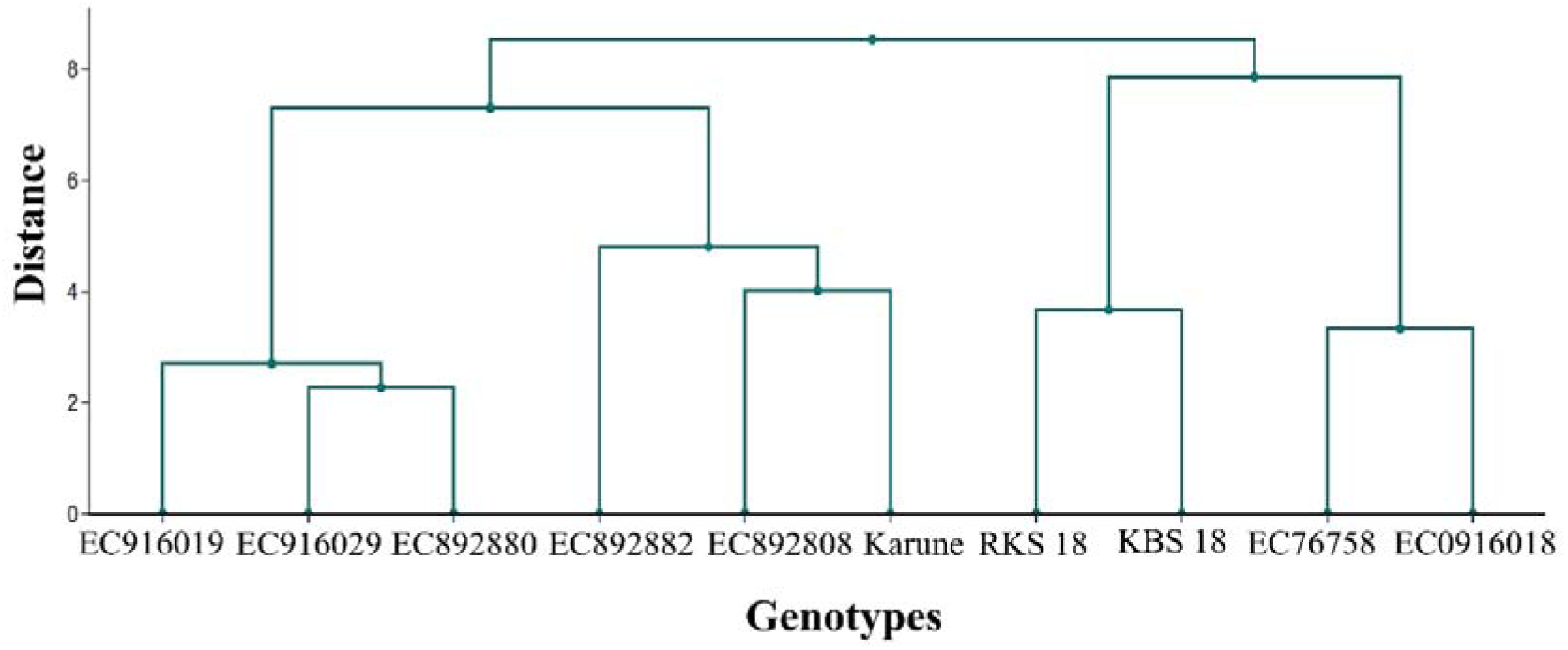
Hierarchical clustering using pheno-morphological traits.

This clustering is similar to Nair *et al*. (2022), who classified soybean accessions into large-seeded (>25g) and small-seeded (<25g) groups, with 456 accessions in the large-seeded and 7397 in the small-seeded group. They created core collections of 1480 small-seeded and 112 large-seeded accessions.

### 3.3. Days taken to achieve the reproductive stages by vegetable and grain type soybean

3.3.1. **R1 stage**

The R1 developmental stage, marked by the appearance of at least one flower on any node, was reached within a range of 37 to 45 days among the experimental soybean lines. The vegetable type line EC892880 was the fastest, reaching R1 in 37 days, while the grain type EC76758 took the longest, with 45 days. Other varieties, such as EC916019 and RKS- 18, took 38 days, and KBS-23, EC0916018, and EC916029 reached R1 in 41 days (**Table 4**).

**Table 4:** Days taken to attain reproductive stages from R1 to R7.

| Sl. No. | Soybean line | Days taken to attain reproductive stages |  |  |  |  |  |  |
| --- | --- | --- | --- | --- | --- | --- | --- | --- |
|  |  | R1 | R2 | R3 | R4 | R5 | R6 | R7 |
| 1 | EC916019 | 38.00 ± 1.00 | 45.00 ± 1.60 | 51.00 ± 0.21 | 63.00 ± 0.30 | 74.00 ± 1.02 | 85.00 ± 0.60 | 92.00 ± 0.10 |
| 2 | EC916029 | 41.00 ± 1.30 | 47.00 ± 1.50 | 55.10 ± 0.31 | 60.10 ± 0.30 | 69.00 ± 1.00 | 79.00 ± 0.40 | 89.00 ± 0.30 |
| 3 | EC892880 | 37.00 ± 1.30 | 42.00 ± 0.30 | 48.00 ± 0.14 | 57.00 ± 0.29 | 67.00 ± 0.68 | 77.00 ± 0.14 | 87.00 ± 0.60 |
| 4 | EC892882 | 40.00 ± 0.50 | 46.00 ± 0.24 | 54.00 ± 0.23 | 61.00 ± 0.31 | 72.00 ± 0.32 | 89.00 ± 0.17 | 95.00 ± 0.50 |
| 5 | EC892808 | 44.00 ± 1.70 | 49.00 ± 0.12 | 53.00 ± 0.34 | 63.00 ± 0.79 | 75.00 ± 0.40 | 84.00 ± 1.70 | 91.00 ± 0.80 |
| 6 | Karunae | 42.00 ± 0.24 | 48.00 ± 0.30 | 52.00 ± 1.60 | 59.00 ± 1.70 | 64.00 ± 0.66 | 70.00 ± 1.07 | 84.00 ± 0.40 |
| 7 | RKS 18 | 38.00 ± 0.42 | 56.00 ± 0.39 | 70.00 ± 0.60 | 81.00 ± 1.10 | 93.00 ± 0.56 | 101.00 ± 0.50 | 112.00 ± 0.10 |
| 8 | KBS 23 | 41.00 ± 0.18 | 52.00 ± 0.79 | 62.00 ± 0.75 | 65.00 ± 0.45 | 75.00 ± 0.90 | 87.00 ± 0.39 | 97.00 ± 1.10 |
| 9 | EC 76758 | 45.00 ± 0.69 | 51.00 ± 0.89 | 63.00 ± 0.40 | 68.00 ± 0.68 | 78.00 ± 0.74 | 89.00 ± 1.07 | 94.00 ± 0.70 |
| 10 | EC 0916018 | 41.00 ± 0.79 | 52.70 ± 0.40 | 64.10 ± 0.20 | 74.00 ± 0.71 | 86.00 ± 0.45 | 96.00 ± 0.10 | 109.00 ± 0.60 |
| Mean |  | 40.70 | 48.87 | 57.22 | 65.11 | 75.30 | 85.70 | 95.00 |
| Minimum |  | 37.00 | 42.00 | 48.00 | 57.00 | 64.00 | 70.00 | 84.00 |
| Maximum |  | 45.00 | 56.00 | 70.00 | 81.00 | 93.00 | 101.00 | 112.00 |
| S. E(m) |  | 1.70 | 1.92 | 0.80 | 2.41 | 2.15 | 0.96 | 0.74 |
| CD (5%) |  | 1.89 | 2.01 | 1.74 | 2.78 | 1.97 | 2.45 | 1.46 |
| CV (Big) |  | 1.46 | 0.93 | 0.84 | 0.91 | 0.82 | 0.85 | 0.78 |

**Note:** CD- Critical difference, SD- Standard deviation, CV- Critical variance, S.E(m)- Standard error mean, R1/2/3/4/5/6/7- Reproductive stage 1/2/3/4/5/6/7

#### 3.3.2. R2 stage

The R2 stage, marked by full flowering in all plants, was reached between 42.00 to 56.00 days across different lines in the study. Significant differences were observed in the time required to reach this stage. The vegetable type cultivar EC892880 took the least time, 42.00 days, while the grain type line RKS-18 took the longest at 56.00 days. The four grain-type check varieties also required the longest duration, ranging from 52.70 to 56.00 days (**Table 4**).

#### 3.3.3. R3 stage

The study observed significant variation in the time taken for different lines to reach the R3 stage of pod development. The vegetable-type EC892880 reached R3 in the shortest time, 48 days, while the grain-type RKS-18 took the longest at 70 days. The vegetable-type check variety, Karune, reached the R3 stage in 52 days, highlighting the variation in time to reach this developmental stage across different types (**Table 4**).

#### 3.3.4. R4 stage

The time to reach the R4 stage, where pod growth ceases, varied significantly among different lines in the field study. The duration ranged from 57 to 81 days. The vegetable-type EC892880 reached R4 the fastest in 57 days, while the grain-type RKS-18 took the longest at 81 days, followed by EC0916018 at 74 days. The vegetable-type check variety, Karune, reached R4 in 59 days (**Table 4**).

#### 3.3.5. R5 stage

The R5 stage, marking the commencement of seed development, showed significant variation in the time required to reach it across different field-grown lines. The vegetable-type check variety, Karune, took the least time (64.00 days), followed by EC892880 (67.00 days). In contrast, the grain-type line EC0916018 took the longest (86.00 days) to reach the R5 stage. Among the lines analyzed, the vegetable type was the first to reach this stage (**Table 4**).

#### 3.3.6. R6 stage

The crop reached the R6 stage after varying durations, ranging from 70 to 101 days. The vegetable-type Karune reached R6 the fastest in 70 days, followed by EC892880 at 77 days. In contrast, the grain-type lines took longer, with RKS-18 taking 101 days and EC0916018 taking 96 days, indicating that grain-type lines required more time to reach this stage (**Table 4**).

#### 3.3.7. R7 stage

The crop reaches the R6 stage when the entire seed population has developed, and the R7 stage is reached after 84 to 112 days, depending on the line. Karune, a vegetable-type check, reached R7 in the shortest time (84 days), while RKS-18, a grain-type line, took the longest (112 days) (**Table 4**).

William *et al*. (2015) found that vegetable soybeans reach the R6 stage 9.5 to 17.9 days earlier than grain types, leading to a shorter growing season. This earlier maturation impacts agronomic practices such as postharvest management, weed control, and the use of cover crops or double plantings.

### 3.4. Genetic variability studies on ten soybean genotypes

The present study focused on quantifying the genetic diversity among 10 soybean genotypes obtained from the Soybean Scheme at UAS, GKVK, Bangalore, with an emphasis on growth and yield-related traits. A total of 11 characteristics were evaluated, and the analysis revealed significant genotypic variations for all traits, indicating a high level of genetic diversity among the genotypes under study. This diversity is essential for improving crop species to meet challenges such as biotic and abiotic stress, as well as for enhancing yield and adaptability (Bhandari *et al*., 2017).

To assess the genetic variability, the phenotypic coefficient of variation (PCV) and genotypic coefficient of variation (GCV) were calculated for each trait (Fischer, 1930). The traits of test weight, number of pods per plant, and plant height at the R7 stage showed high PCV and GCV, suggesting that both genetic and environmental factors contribute to their variation. On the other hand, traits such as the number of pods per cluster, number of seeds per pod, and days to 50% flowering exhibited lower values for both PCV and GCV, indicating that these traits are less influenced by environmental factors and have a stronger genetic basis (**Table 5**).

**Table 5:** Estimation of genetic parameters in eleven characters of ten genotypes in vegetable type and grain type soybean.

| Traits | Mean | Range |  | PCV | GCV | h<sup>2</sup> | GAM | GA | EV |
| --- | --- | --- | --- | --- | --- | --- | --- | --- | --- |
|  |  | Minimum | Maximum |  |  |  |  |  |  |
| **DPS** | 68.482 | 66.06 | 71.00 | 3.54 | 3.43 | 93.64 | 6.84 | 4.54 | 0.11 |
| **DTRR7** | 78.382 | 77.26 | 80.40 | 4.97 | 1.14 | 81.41 | 2.12 | 1.65 | 2.35 |
| **DTF** | 42.48 | 40.00 | 45.00 | 5.44 | 5.43 | 99.89 | 11.15 | 4.74 | 0.03 |
| **TW** | 20.38 | 14.49 | 24.72 | 24.65 | 24.64 | 99.74 | 50.75 | 11.46 | 0.01 |
| **NBPP** | 12.4 | 10.66 | 14.60 | 17.75 | 15.11 | 67.34 | 2.09 | 2.30 | 3.01 |
| **NPPC** | 6.34 | 5.26 | 6.93 | 9.62 | 4.89 | 50.47 | 8.83 | 5.60 | 5.85 |
| **NPPP** | 31.82 | 23.86 | 39.60 | 32.34 | 32.09 | 98.58 | 65.60 | 21.73 | 0.78 |
| **NSPP** | 2.63 | 2.18 | 2.98 | 6.87 | 1.61 | 54.44 | 7.74 | 2.10 | 4.03 |
| **PHR7** | 49.73 | 31.07 | 65.24 | 29.65 | 29.50 | 98.57 | 60.45 | 29.94 | 0.25 |
| **PL** | 5.27 | 4.69 | 5.63 | 7.29 | 6.48 | 79.36 | 11.87 | 6.16 | 1.03 |
| **SS** | 8.27 | 6.03 | 10.64 | 19.41 | 18.80 | 93.11 | 37.50 | 3.10 | 2.06 |

**Note:** **PCV**- Phenotypic coefficient of variation, **GCV**- Genotypic coefficient of variation, **h<sup>2</sup>**- Heritability (broad sense), **GAM**- Genetic advance as mean, **GA**- Genetic advance, **EV**- Environmental variance, **DPS**: Days for pod setting; **DTRR7**: Days Taken to Reach R7 stage; **DTF**: Days To 50% Flowering; **TW**: Test Weight; **NBPP**: Number of Branches Per Plant; **NPPC**: Number of Pods Per Cluster; **NPPP**: Number of Pods Per Plant; **NSPP**: Number of Seeds Per Pod; **PHR7**; Plant Height at R7 stage; **PL**: Pod Length; **SS**: Seed Size

Heritability estimates in the broad sense were calculated for each trait, and the results showed that traits like days to 50% flowering, test weight, and the number of pods per plant had high heritability (above 90%). High heritability indicates that these traits are largely controlled by genetic factors and are likely to respond well to selection in breeding programs (Das *et al*., 2024). On the other hand, traits such as the number of pods per cluster and number of seeds per pod had lower heritability, suggesting that environmental factors may play a more significant role in their expression (**Table 5**).

The genetic advance as mean (GAM) (Johnson *et al*., 1955), which indicates the expected improvement in a trait through selection, was high for the number of pods per plant (65.6%), plant height at the R7 stage (60.45%), and test weight (50.75%). These traits are promising for genetic improvement due to their high genetic variability and high GAM. Traits like pod length and days to 50% flowering showed moderate GAM, while traits such as the number of pods per cluster, number of seeds per pod, and days for pod setting exhibited low GAM, suggesting limited potential for improvement through selection (**Table 5**).

In terms of genetic advance (GA), the highest values were observed for plant height at the R7 stage (29.94) and the number of pods per plant (21.73), indicating that these traits can be effectively improved through selection. The study concludes that significant genetic diversity exists among the soybean genotypes, particularly for traits like test weight, plant height, and pod number, which are critical for enhancing productivity and stress resistance in breeding programs. These findings underscore the potential for utilizing the genetic variability present in the soybean genotypes to develop improved cultivars (**Table 5**).

In the study, it was observed that for most traits, the phenotypic coefficient of variation (PCV) was higher than the genotypic coefficient of variation (GCV), indicating that environmental factors influence trait expression. However, in traits such as days for pod setting, days to 50% flowering, test weight, number of pods per plant, and plant height at R7 stage, GCV was nearly equal to PCV, suggesting that genetic factors play a predominant role and environmental impact is minimal. In contrast, traits like days to reach R7 stage, number of pods per cluster, and number of seeds per pod exhibited a larger difference between PCV and GCV, indicating a significant environmental influence on variation (**Table 5**).

High heritability and genetic advance were observed for traits like number of pods per plant, plant height at R7 stage, and test weight, suggesting additive gene action and the possibility of improvement through simple selection methods like pure line or mass selection. However, traits like number of pods per cluster and number of seeds per pod showed moderate heritability and higher genetic advance, indicating that environmental factors still play a role, and selection should focus on recombination breeding in later generations (**Table 5**).

The study’s results are consistent with those of Khanande *et al*. (2016), where similar patterns of high heritability and low environmental influence were reported for traits like pod length, 100 beans weight, and fresh pod weight. Traits with low environmental impact and high genetic variation are more likely to respond well to phenotypic selection, and a high GCV, combined with heritability and genetic advance, suggests promising prospects for breeding improvements.

### 3.5. Principal component analysis

Principal Component Analysis (PCA) is widely used in agricultural studies to reduce data dimensionality. Rotaru *et al*. (2012) highlighted its benefits for understanding genetic variations in phenotypic traits. In a study on 10 soybean genotypes, PCA revealed that the first two principal components (PC1 and PC2) explained 84.5% of the total variation.

The genotypes contributing most to PC1 were KBS-23, EC76758, EC0916018, and RKS 18. Conversely, EC916019, EC916029, EC892880, EC892882, EC892808, and Karune contributed negatively. For PC2, the highest contributors were EC76758 and EC0916018, with negative contributions from EC916019, EC892880, EC892808, RKS-18, and KBS-23 (**Table 6**).

**Table 6:**
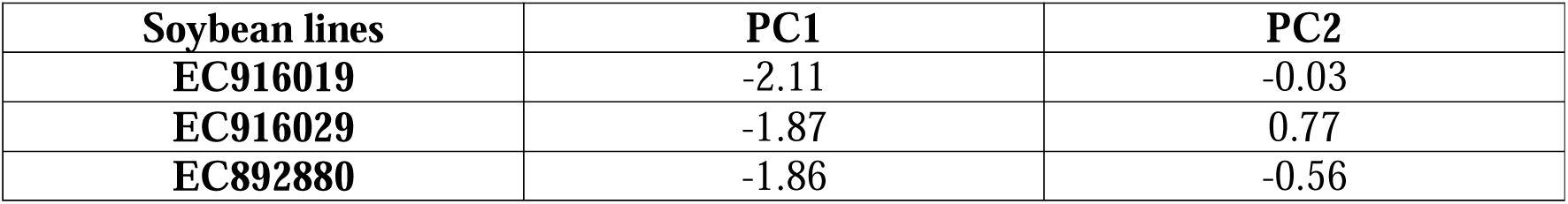
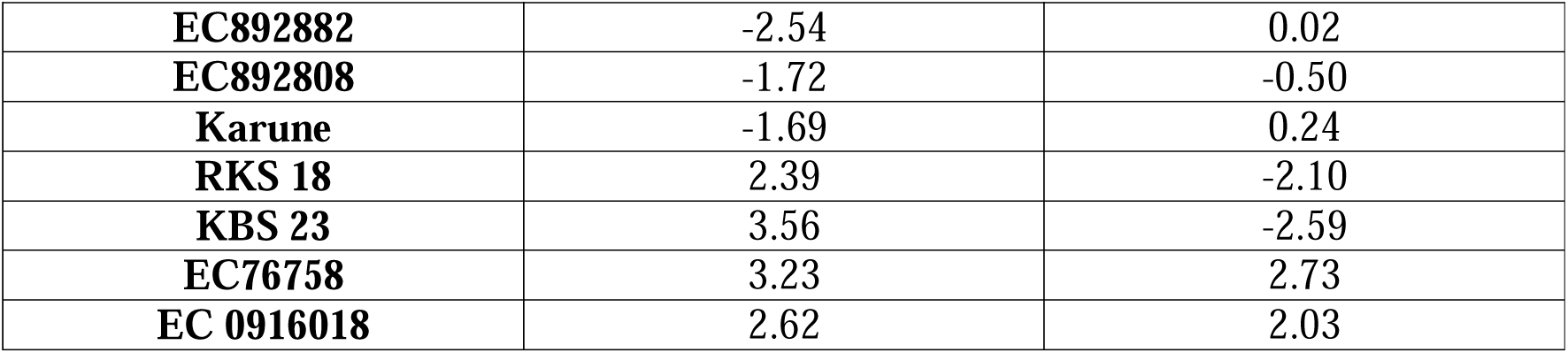
Contribution of morphological traits in soybean to variation in PC1 and PC2.

Key traits contributing to PC1 included plant height at R7 stage, seed size, days for pod setting, number of branches, pod length, number of seeds per pod, number of pods per cluster, and test weight. PC2 was mostly influenced by number of pods per cluster, pod length, days for pod setting, number of pods per plant, test weight, and number of branches, with many traits contributing negatively (**Table 7**).

**Table 7:**
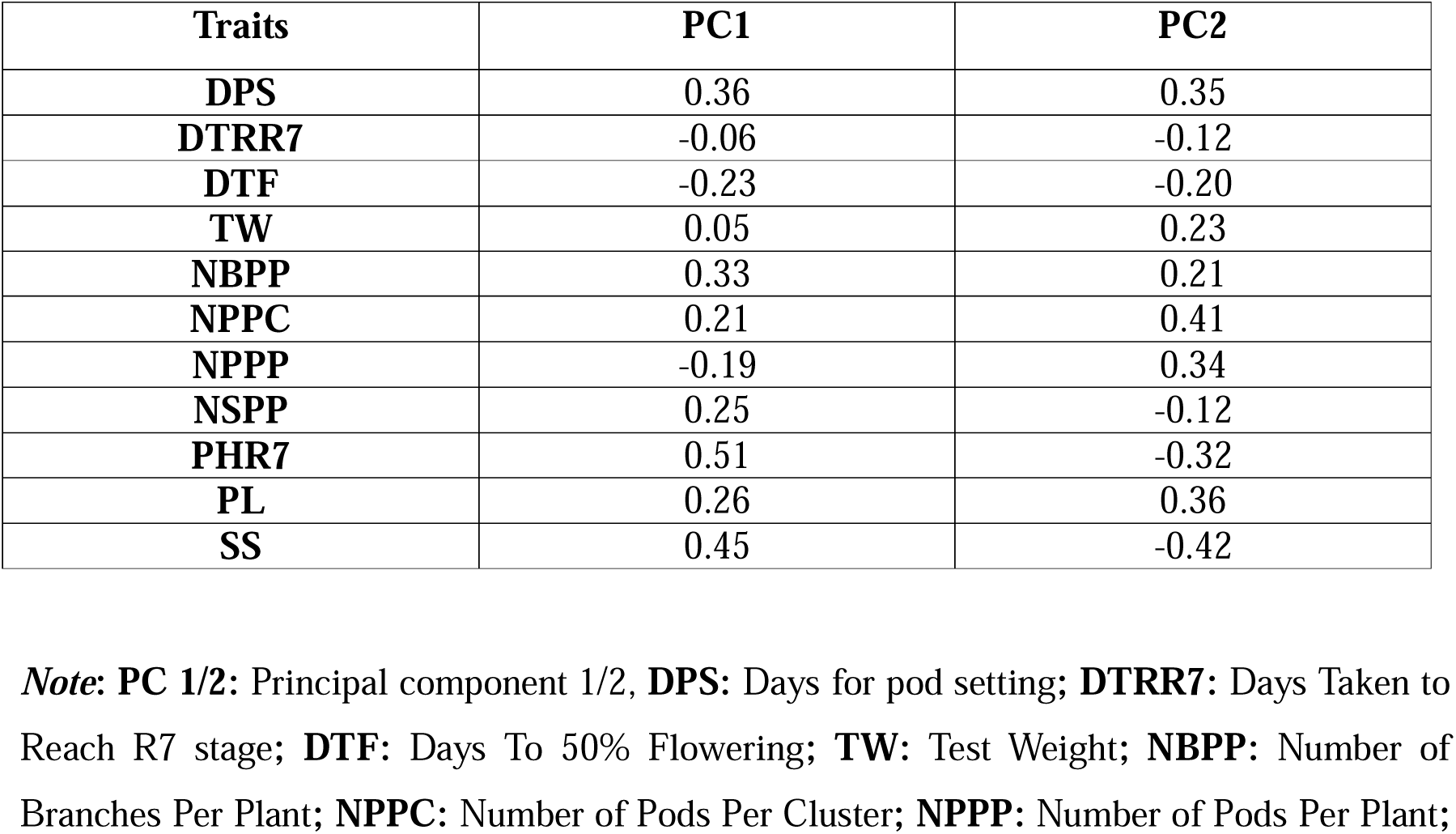
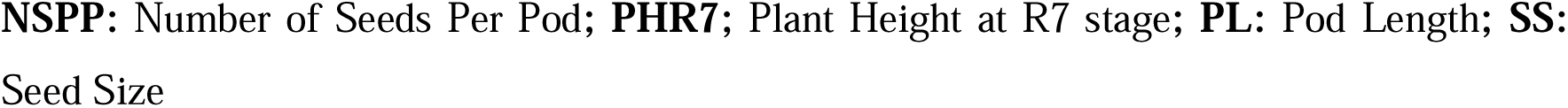
Contribution of soybean genotypes to variation in PC1 and PC2.

| **Traits** | **PC1** | **PC2** |
| --- | --- | --- |
| **DPS** | 0.36 | 0.35 |
| **DTRR7** | -0.06 | -0.12 |
| **DTF** | -0.23 | -0.20 |
| **TW** | 0.05 | 0.23 |
| **NBPP** | 0.33 | 0.21 |
| **NPPC** | 0.21 | 0.41 |
| **NPPP** | -0.19 | 0.34 |
| **NSPP** | 0.25 | -0.12 |
| **PHR7** | 0.51 | -0.32 |
| **PL** | 0.26 | 0.36 |
| **SS** | 0.45 | -0.42 |

**Note: PC 1/2:** Principal component 1/2, **DPS:** Days for pod setting; **DTRR7:** Days Taken to Reach R7 stage; **DTF:** Days To 50% Flowering; **TW:** Test Weight; **NBPP:** Number of Branches Per Plant; **NPPC:** Number of Pods Per Cluster; **NPPP:** Number of Pods Per Plant;
**NSPP:** Number of Seeds Per Pod; **PHR7;** Plant Height at R7 stage; **PL:** Pod Length; **SS:** Seed Size

Traits like plant height at the R7 stage, number of branches per plant, and number of seeds per pod significantly contribute to the first principal component (Dim1), which explains the majority of the phenotypic variation among genotypes. These traits could be prioritized in breeding programs to enhance desirable characteristics. Conversely, traits with a negative contribution, indicated by vectors pointing in the opposite direction along Dim2, should be managed or combined with positively contributing traits for optimal performance (**Table 7**).

The PCA scatter plot showed distinct clustering of genotypes in different quadrants, reflecting their genetic and phenotypic variation. PC1 primarily explained the variation driven by traits like pod length and number of pods per cluster, while PC2, though explaining less variance, was influenced by traits like plant height and days for pod setting. These findings provide insights into key traits for soybean breeding programs aimed at enhancing variability (**Figure 4A**).

**Figure 4:**
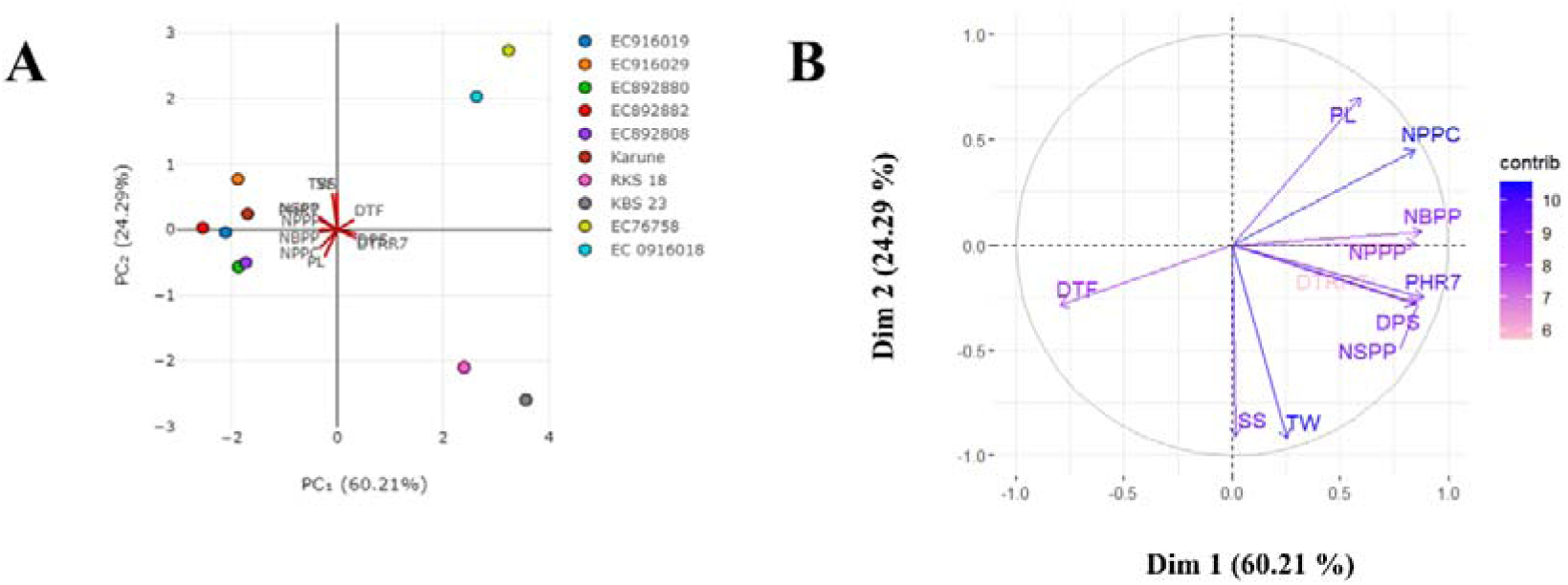
**A.** Scatter plot for pheno-morphological traits showing similarities among genotypes; **B.** Biplot for pheno-morphological traits showing contribution of each trait to principal component

The colour intensity of the vectors reflects each trait’s contribution to the principal components, with darker colours indicating higher contributions. Pod length, number of pods per cluster, and number of branches per plant significantly contribute to Dim1, highlighting their importance in explaining variation along this component. Plant height at the R7 stage, days to pod setting, and number of seeds per pod influence both Dim1 and Dim2. Traits close to each other on the biplot are positively correlated, such as the number of pods per cluster and number of branches per plant. Conversely, traits pointing in opposite directions, like days to 50% flowering and test weight, are negatively correlated (**Figure 4B**).

Out of 11 principal components, 2 components had eigenvalues greater than 1.00, explaining 84.5% of the total variation. The remaining 9 components, with eigenvalues less than 1.00, contributed only 15.5%. Among the traits, PC1 explained 60.21%, and the cumulative variability across PCs 2 to 11 progressively increased, reaching 99.9994%.

Eigenvalues indicate the amount of variance each component explains, with higher values corresponding to greater variance capture. The gradual decline in eigenvalues suggests that the first components are more significant, while the later ones still contribute to understanding the data structure (**Table 8**).

**Table 8:** Principal components (PCs) analysis for metric traits in soybean genotypes.

| Characters | Principal component | Eigen values | Percentage of variance | Cumulative % of variance |
| --- | --- | --- | --- | --- |
| **DPS** | PC1 | 6.62350 | 60.2136 | 60.2136 |
| **DTRR7** | PC2 | 2.67160 | 24.2872 | 84.5008 |
| **DTF** | PC3 | 0.74680 | 6.78930 | 91.2901 |
| **TW** | PC4 | 0.43610 | 3.96450 | 95.2546 |
| **NBPP** | PC5 | 0.24510 | 2.22790 | 97.4825 |
| **NPPC** | PC6 | 0.14320 | 1.30230 | 98.7847 |
| **NPPP** | PC7 | 0.07620 | 0.69350 | 99.4783 |
| **NSPP** | PC8 | 0.04930 | 0.44810 | 99.9264 |
| **PHR7** | PC9 | 0.00950 | 0.07359 | 99.9810 |
| **PL** | PC10 | 0.000288 | 0.00026 | 99.9901 |
| **SS** | PC11 | 0.000080 | 0.000043 | 99.9994 |

**Note:** **DPS:** Days for pod setting; **DTRR7:** Days Taken to Reach R7 stage; **DTF:** Days To 50% Flowering; **TW:** Test Weight; **NBPP:** Number of Branches Per Plant; **NPPC:** Number of Pods Per Cluster; **NPPP:** Number of Pods Per Plant; **NSPP:** Number of Seeds Per Pod; **PHR7:** Plant Height at R7 stage; **PL:** Pod Length; **SS:** Seed Size

The scree plot from PCA shows the distribution of eigenvalues across 11 principal components (PCs) and their cumulative variance. PC1, with the highest eigenvalue of 6.6235, explains the majority of the variance, followed by PC2 with an eigenvalue of 2.6716. Together, these two components account for around 84.5% of the total variance, as seen in the steep rise of the cumulative variance curve. The remaining components (PC3 to PC11) have eigenvalues below 1, contributing minimally to the overall variance. Thus, PC1 and PC2 are the key components for further analysis, while the others are less significant (**Figure 5**).

**Figure 5:**
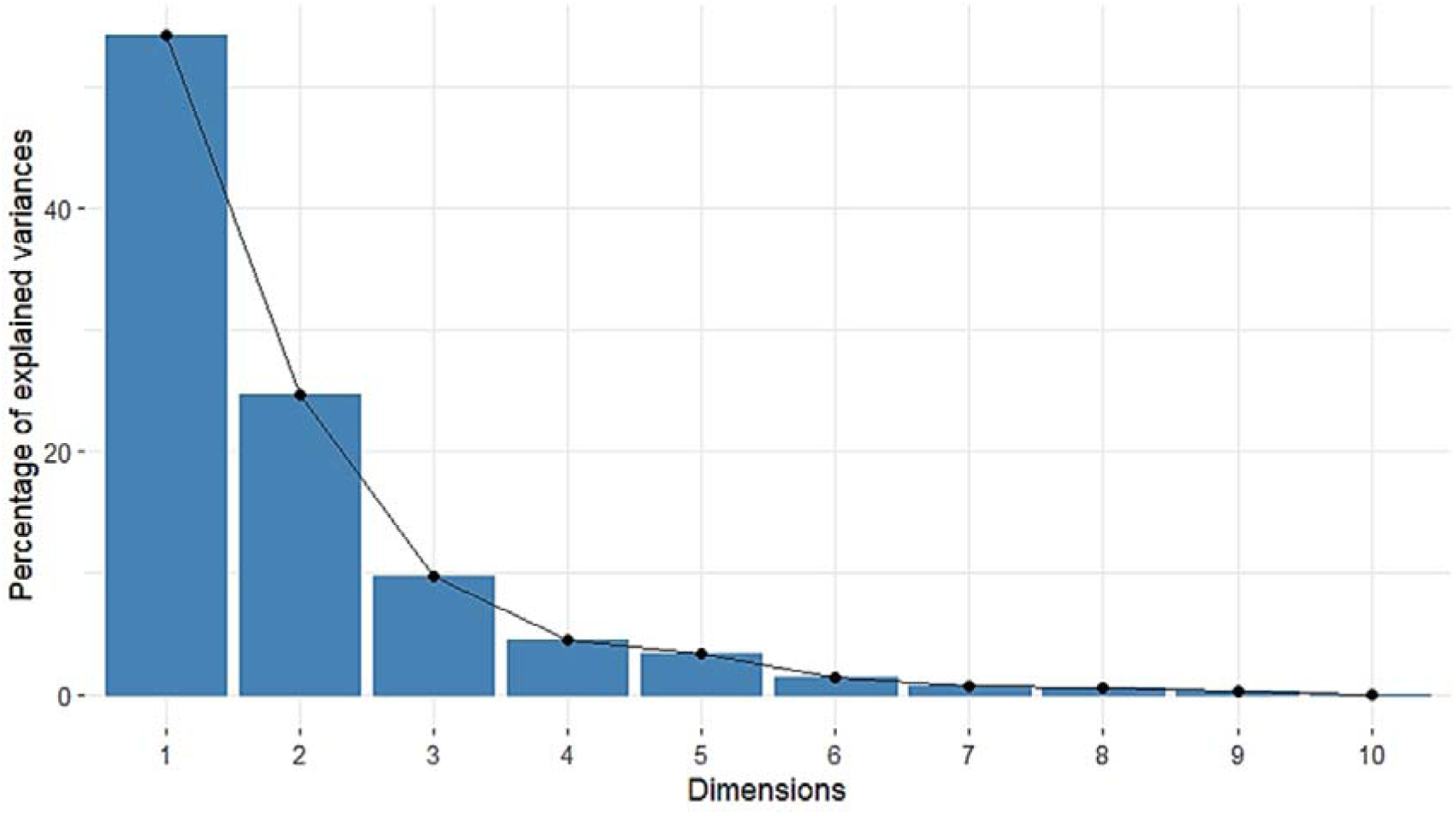
Scree plot for pheno-morphological traits showing contribution of each principal component.

Dunna *et al*. (2023) evaluated 33 vegetable soybean genotypes for 14 quantitative traits during the Kharif season of 2021, finding significant variation. Principal component analysis (PCA) identified seven components with eigenvalues >1, explaining 81.748% of the total variation. Genotypes EC 915989 and EC 915900 scored high on PC1, indicating superior performance in traits like 100 fresh pod weight and 100 fresh seed weight.

### 3.6. Diversity analysis and grouping based on morphological data

Soybean genetic divergence was assessed using data from 11 traits across 10 genotypes, with clustering performed using Tocher’s method and Mahalanobis D² distance (Rao, 1952). The genotypes were grouped into three clusters: Cluster 1 and Cluster 2 each contained 2 genotypes, while Cluster 3 had 6 genotypes (**Table 9**). According to Singh’s statistics (1981), the trait contributing most to genetic diversity was the days to reach the R7 stage (18.163%), followed by the number of pods per plant (7.6148%) (**Table 10**).

**Table 9:**
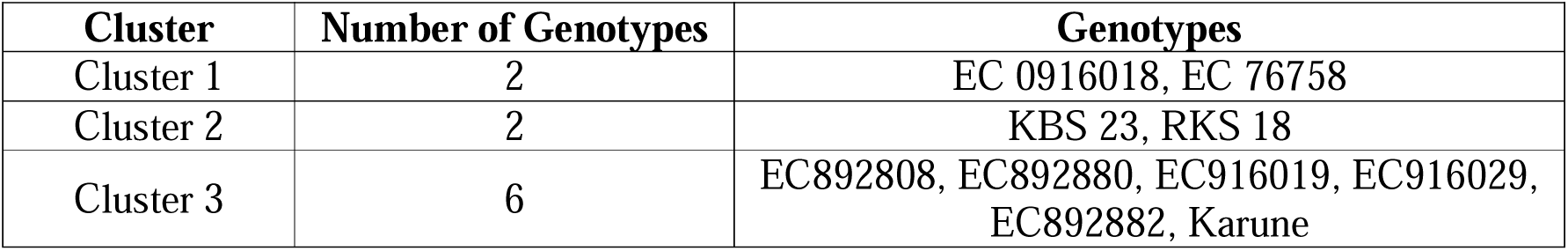
Grouping of soybean genotypes into three clusters based on various morphological traits.

| Cluster | Number of Genotypes | Genotypes |
| --- | --- | --- |
| Cluster 1 | 2 | EC 0916018, EC 76758 |
| Cluster 2 | 2 | KBS 23, RKS 18 |
| Cluster 3 | 6 | EC892808, EC892880, EC916019, EC916029, EC892882, Karune |

**Table 10:** Percentage contribution of all the traits for diversity in soybean genotypes.

| Traits | Singh statistic | % contribution for diversity |
| --- | --- | --- |
| DPS | 0.356 | -1.037 |
| DTRR7 | -0.01037 | 18.163 |
| DTF | 0.027778 | 3.1111 |
| TW | 0.013007 | 1.3567 |
| NBPP | -0.02274 | -10.2815 |
| NPPC | 0.007259 | -3.2 |
| NPPP | 0.169926 | 7.6148 |
| NSPP | -0.03822 | -1.2296 |
| PHR7 | 0.298778 | -14.4622 |
| PL | 0.026237 | 1.7541 |
| SS | -0.0961 | -0.1704 |

**Note:** **DPS:** Days for pod setting; **DTRR7:** Days Taken to Reach R7 stage; **DTF:** Days To 50% Flowering; **TW:** Test Weight; **NBPP:** Number of Branches Per Plant; **NPPC:** Number of Pods Per Cluster; **NPPP:** Number of Pods Per Plant; **NSPP:** Number of Seeds Per Pod; **PHR7:** Plant Height at R7 stage; **PL:** Pod Length; **SS:** Seed Size

The maximum genetic distance was found between cluster II and III (D^2^ = 28292.49), followed by cluster I and III (D^2^ = 9650.051), then by cluster I and II (D^2^ = 7594.65) (**Figure 6**).

**Figure 6:**
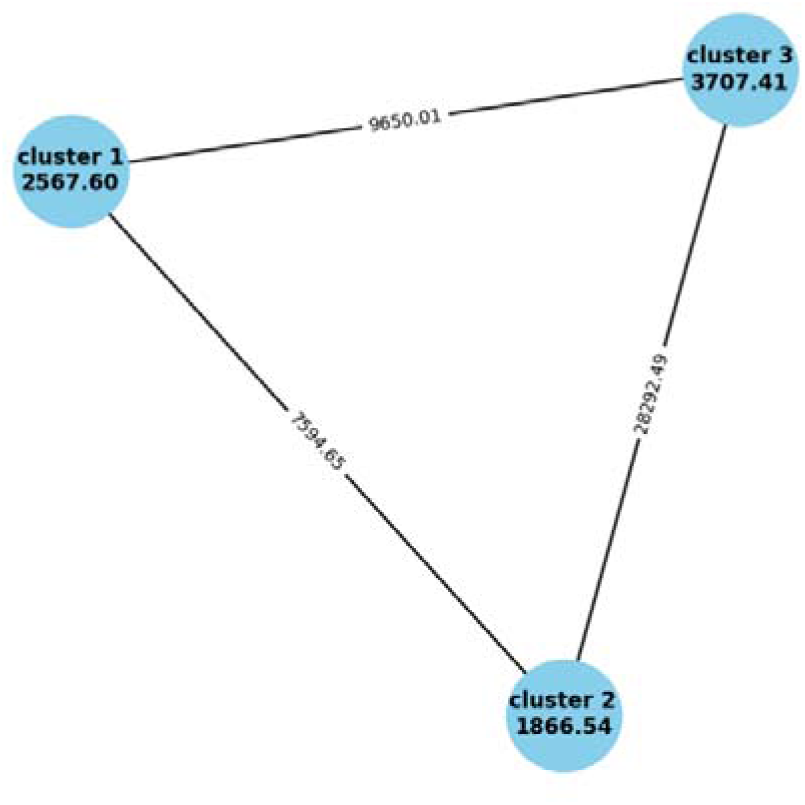
Clustering of genotypes using Tocher’s method and Mahalanobis D^2^ distance with edge weights based on pheno-morphological traits.

In the study by Shilpashree *et al*. (2021), vegetable soybean divergence was assessed using 18 traits across 28 genotypes, with clusters formed by Trocher’s method. The genotypes were grouped into eight clusters, with Cluster II containing the most genotypes (8), followed by Cluster V and Cluster I (each with 4 genotypes). Clusters III and IV had 3 genotypes, while Clusters VI, VII, and VIII had 2 each. The greatest genetic distance (D2 = 51,828.79) was observed between Clusters VIII and I, with genotypes GM-6 and GM-27 from Cluster VIII, and GM-10, GM-18, GM-20, and GM-25 from Cluster I showing the highest genetic distance.

## 4. Conclusion

The study involving vegetable and grain soybean genotypes revealed significant differences in traits such as plant height, pod characteristics, seed size, and flowering times. Vegetable-type soybeans tended to flower earlier and mature faster with generally greater plant heights than grain types. Genotype EC892882 exhibited the earliest flowering time and longer pods, while the tallest plants were observed in genotype EC916029. Grain-type soybeans produced larger and heavier seeds compared to vegetable types, with EC76758 showing the largest seed size. Hierarchical clustering identified three major groups: one characterized by vegetable-type soybeans with distinct traits such as pubescence density and corolla color, another by grain-type soybeans with evident grouping, and a third cluster highlighting intermediate traits.

Correlation analysis revealed positive associations among yield-related traits such as plant height, seed size, and the number of branches, while days to pod setting showed a negative correlation. Traits like plant height and seed size were identified as important contributors to phenotypic variation, emphasizing both genetic control and environmental influence. High heritability was noted for traits such as plant height and flowering time, suggesting their potential for targeted breeding programs. However, traits such as pod length and pod count per cluster were found to be more environmentally influenced.

An analysis of genetic diversity using Mahalanobis D2 distance confirmed three major clusters, with significant genetic distances between vegetable and grain types, showcasing the potential for breeding improvements. This diversity, coupled with the strong associations of traits, highlights opportunities for developing soybean genotypes optimized for diverse growing conditions and market demands. The findings also suggest that vegetable soybeans can be improved for faster growth and shorter growing seasons, leveraging their earlier flowering maturity and taller plant heights. The results underscore the breeding potential of genotypes such as EC892882 for early flowering and EC76758 for large seed size, offering considerable promise for enhancing soybean performance and adaptability.

## 7. Author Contributions

The authors confirm contribution to the paper as follows: Madhumita Saha: study conception, design, analysis, interpretation of results, preparation of manuscript and writing; Mohan Chavan: interpretation of results, preparation of manuscript, revision, reviewing and editing; Onkarappa T: revision, reviewing and editing; Ravikumar R L: reviewing and editing; Uddalak Das: reviewing and editing. All authors have read and agreed to the published version of the manuscript.

## 8. Data Availability

The datasets analysed during the current study available from the corresponding author on reasonable request. No third part data was used in this paper. All are original and included in this MS.

## 9. Ethics declarations

### Ethics approval and consent to participate

The plants used in the study is owned to the breeder who generated them who is Dr. Onkarappa T. (one of the authors) who is the Head of the Soybean Scheme at ZARS (Zonal Agricultural Research Station) during the time the experiments and the study were taken place. Therefore, the collection of the plants used in the study complies with local or national guidelines with no need for further affirmation. The material used in this study was based upon work supported by the Soybean Scheme, ZARS, UAS Bangalore-560065. Any opinions, findings, conclusions, or recommendations expressed in this publication are those of the author(s) and do not necessarily reflect the view of the Soybean Scheme, ZARS, UAS Bangalore.

## Acknowledgement

None

